# EPISOMAL VECTORS FOR STABLE PRODUCTION OF RECOMBINANT PROTEINS AND ENGINEERED ANTIBODIES

**DOI:** 10.1101/2024.01.03.574076

**Authors:** Ian Fallahee, Daniel Hawiger

## Abstract

There is tremendous interest in the production of recombinant proteins, particularly bispecific antibodies and antibody-drug conjugates for research and therapeutic use. Here, we demonstrate a highly versatile plasmid system that allows rapid generation of stable Expi293 cell pools by episomal retention of transfected DNA. By linking protein expression to puromycin resistance though an attenuated internal ribosome entry site, we achieve stable cell pools producing proteins of interest. In addition, split intein-split puromycin-mediated selection of two separate protein expression cassettes allows the stable production of bispecific antibody-like molecules or antibodies with distinct C-terminal heavy chain modifications, such as an antigen on one chain and a sortase tag on the other chain. We also use this novel expression system to generate stable Expi293 cell pools that secrete sortase A Δ59 variant Srt4M. Using these reagents, we prepared a site-specific drug-to-antibody ratio of 1 antibody-siRNA conjugate. We anticipate the simple, robust, and rapid stable protein expression systems described here being useful for a wide variety of applications.

## INTRODUCTION

Recombinant engineered antibodies have become commonplace in modern medicine^1,2^ and are typically produced by transient transfection for research purposes. For large scale biotherapeutic manufacturing, Chinese Hamster Ovary (CHO) cells are dominant but there are increasing examples of Human Embryonic Kidney (HEK) 293-based cells being used for production of FDA-approved therapeutics^3^. While production in CHO cells is suitable for standard antibodies with limited glycosylation^4,5^, more complex molecules, such as protein-Fc fusions, may contain non-human carbohydrates such as galactose-α-1,3-galactose^6^ and N-glycolylneuraminic acid^7^. Therefore, this warrants the further development of human-derived expression systems, which eliminate the risk of specific allergic responses and anaphylactic reactions stemming from the immunogenicity of these non-human carbohydrates^6–8^. HEK293-based cell lines are widely used in protein production for research^9^ and the availability of current good manufacturing practices (cGMP)-banked Expi293F lines^10^ affords the potential to transition to commercial production. Therefore, methods that simplify and accelerate the stable production of proteins in HEK293-based cells are of significant interest.

Stable protein production typically involves integration of the transgene of interest into the cell line’s genome, a random and low probability event following transfection of linearized DNA^11^. More efficient integration can be achieved by lentivirus transduction^12^ or by co-transfection of the transgene of interest (flanked by inverted terminal repeats) with a transposase plasmid^13–16^. Here, we demonstrate a simpler approach using episomal vectors that allows rapid generation of stably-transfected Expi293 cell pools without the need for lentiviral transduction or exogenous transposons.

Scaffold/matrix attachment region (S/MAR) elements and replication initiation (IR) elements have previously been used in vectors for recombinant protein production^17–19^ or for gene therapy of human cells^20^. S/MAR elements contain sequences that serve as binding sites for proteins such as scaffold attachment factor A, which tethers these elements to the nuclear matrix^21^. IR elements act as a mammalian origin of replication, initiating DNA replication with each cell division^22^. Depending on the particular arrangement of S/MAR and IR elements, plasmids can be integrated and amplified in the genome as a result of double stranded (ds) DNA breaks^23^ or they can be maintained episomally without integration or gene amplification^20^. To avoid disruption of the expressed gene of interest by random DNA breaks^24^, we chose an episomal design of S/MAR and IR elements previously established by Stavrou et al.^20,25^. This design was shown to allow long-term replication of the introduced plasmid once per cell cycle in human hematopoietic progenitor cells^20^.

As episomal vectors do not undergo gene amplification, we chose an alternative way to increase protein production. Within these vectors we coupled protein expression to puromycin resistance (PuroR) by an attenuated internal ribosome entry site (IRES)^26^. An attenuated IRES reduces the expression of the PuroR protein relative to the protein of interest, allowing for increased selection stringency that results in higher producing cells^26,27^. This linked PuroR approach is additionally beneficial for long-term pool stability as the coupling of the expression of protein and selectable marker to a single promoter reduces the chances of selective silencing of the protein of interest^27,28^. Here, we show that episomal retention combined with protein-linked attenuated expression of PuroR allows rapid generation of stable Expi293 pools producing recombinant proteins of interest.

One of the recombinant proteins we produced by secretion into supernatants was Sortase A Δ59 variant P94S/D160N/D165A/K196T (Srt4M)^29,30^, a highly efficient bioconjugation enzyme^29^. Secretion of recombinant proteins in mammalian cells, especially those not naturally secreted, is a non-trivial task^31^. Attempts have been made to generate synthetic signal peptides based on machine-learning^31–33^ but these methods are not yet fully reliable. Therefore, rather than use a synthetic signal peptide, we took inspiration from the findings of Güler-Gane et al.^31^ by using natural mammalian signal peptides with predicted cleavage that matches at least the +1/+2 residues of the protein of interest. This strategy revealed several novel signal peptides that enable secretion of Srt4M in mammalian cells.

The main class of recombinant proteins we focused on was engineered antibodies. Using the attenuated PuroR expression approach, we demonstrated stable expression of engineered antibodies with C-terminal antigens or tags. C-terminally modified antibodies are typically used for sensitive immunological studies in mice^34–36^ and cancer immunotherapy in humans^37,38^. However, when it comes to producing bispecific antibody-like molecules or antibodies with distinct C-terminal modifications on each chain, at least two different expression cassettes are required. By splitting the PuroR protein using a recently described split-intein system^39^, we efficiently selected for two separate expression cassettes. Specifically, we made use of the SiMPl split-intein split-puromycin system, described by Palanisamy et al.^39^, to produce stable Expi293 cell pools expressing a bispecific antibody-protein-Fc fusion and antibodies with multiple distinct heavy chain C-terminal tags, including a single sortase conjugation site. Ultimately, we demonstrated the applicability of these new approaches by preparing a site-specific drug-to-antibody ratio of 1 (DAR1) antibody-siRNA conjugate by Srt4M-mediated ligation and click chemistry^40^.

## RESULTS

### Secreted expression of sortase A Δ59 variant Srt4M in Expi293

Sortase A is used as a versatile bioconjugation reagent for site-specific attachment of oligoglycine-containing molecules to an LPXTG amino acid sequence (sortase tag) present within another protein in a process referred to as sortagging^41,42^. As a protein of bacterial origin, expression of sortase A is typically achieved in E. coli; however, production in typical strains inevitably results in lipopolysaccharide (LPS, endotoxin) contamination, complicating further research and clinical applications^43,44^. Therefore, we aimed to express sortase A Δ59 variant Srt4M^29,30^, recently described to reach maximal ligation in as little as 15 minutes^29^, in mammalian cells to eliminate endotoxin and therefore extend its use to LPS-free applications.

To express Srt4M, we designed an episomal vector with a single attenuated IRES^26,27^ from encephalomyocarditis virus (EMCV) (bicistronic cassette) (Fig. 1A). This bicistronic cassette contains an immediate early human CMV enhancer and promoter leading to expression of Srt4M with C-terminal His_6_ tag as the first cistron and linked by an attenuated IRES to an enhanced green fluorescent protein-self cleaving porcine teschovirus 2A peptide^45^-puromycin N-acetyltransferase (EGFP-P2A-PuroR) selection cistron further described in Methods. The self-cleaving P2A peptide allows EGFP and PuroR to separate during translation and function as independent proteins^46^. The complete expression vector also contains an S/MAR from human β-interferon^47^ before the SV40 poly A signal, and the minimal β-globin IR, G5^48^, outside of the main expression cassette. Episomal elements, in combination with linked attenuated IRES-mediated translation of the EGFP-P2A-PuroR cistron, allow the generation of stable Expi293 pools expressing a protein of interest by puromycin selection. It is also possible to monitor relative protein expression by EGFP. We included a vector backbone map listing restriction enzymes in Fig. S1A with expanded cloning details in Methods.

**Figure 1.**
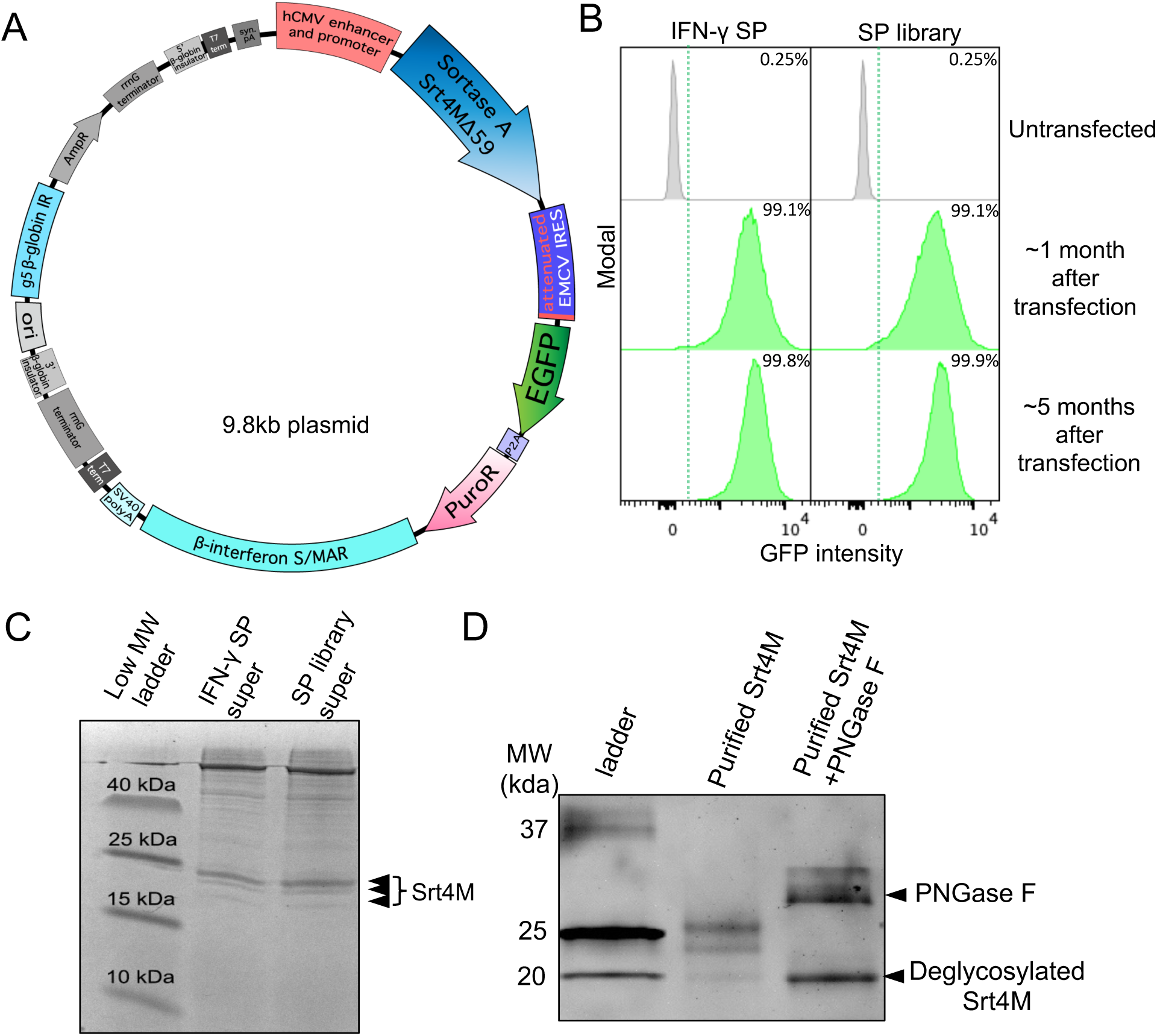
Specific signal peptides allow secretion of Srt4M in Expi293 cells. **A.** Bicistronic Sortase A Srt4M Δ59 (Srt4M) plasmid map. Elements include, as indicated: human CMV (hCMV) promoter and enhancer, Srt4M, attenuated EMCV IRES, EGFP, P2A self-cleaving peptide, PuroR, β-interferon S/MAR, SV40 poly A (pA), G5 β-globin replication initiation (IR) **B.** Histograms show GFP expression intensity among untransfected Expi293 cells or cells that were transfected with Srt4M plasmid as in A using either the IFNγ signal peptide (SP) (left panel) or a SP library (right panel) (see text for details) and were then subject to selection with puromycin (see text) for 1 or 5 months as indicated. Percentages denote the frequency of GFP^+^ cells in corresponding cultures as indicated. **C.** Supernatants from stably-transfected Expi293 cell pools from B after 4-5 months of selection were resolved on an SDS-PAGE gel and are shown as indicated. **D.** SDS-PAGE gel showing Srt4M from the SP library pool of stably-transfected Expi293 cells. Nickel column-purified Srt4M with and without treatment with PNGase F deglycosylation is shown as indicated.

Sortase A Δ59^30,49^, a truncated version of the bacterial protein, has been engineered for secretion in bacteria^50^, but to our knowledge has not been previously secreted in mammalian cells. Secretion of proteins in mammalian cells is a non-trivial task and attempts have been made to find “universal” signal peptides such as secrecon^31^. Güler-Gane et al. achieved secreted expression of the proteins they studied using secrecon or secrecon with added N-terminal alanines, but found secretion was especially unpredictable for proteins starting with cysteine, proline, tyrosine or glutamine (sortase A Δ59 begins with glutamine). We first attempted to express Srt4M with secrecon and two alanines and were unable to achieve expression (as monitored by GFP) or generate a stable cell pool (data not shown). This is consistent with the “all or nothing” phenomenon described by Güler-Gane et al. where inappropriate signal peptides achieve no expression. Instead of trying secrecon alone or secrecon plus a single alanine as previously described^31^, we explored an alternative approach by using signal peptides from naturally secreted proteins with predicted signal peptide cleavage that exactly matched the +1/+2 amino acids of Srt4M (Gln-Ala).

We first tried the IFN-γ signal peptide from *Ailuropoda melanoleuca* (Giant Panda) as we reasoned that IFN-γ would be expressed highly in its natural context and the predicted +1/+2 amino acids perfectly matched those of Srt4M. Using the IFN-γ signal peptide, we achieved a cell pool expressing Srt4M after about 1 month of selection as indicated by the IRES-linked GFP expression measured by flow cytometry relative to an untransfected control (Fig. 1B, left panel). The general gating strategy is shown in Fig. S1B and applies to all subsequent histograms of GFP expression. In parallel, we also employed a signal peptide library approach using 17 different signal peptides from naturally secreted proteins that all perfectly matched the predicted +1/+2 cleavage (Gln-Ala) (Table S1). A mix of plasmids containing the library of 17 signal peptides for expression of Srt4M was simultaneously transfected into Expi293 cells followed by selection, resulting in a viable GFP+ cell pool after the first month (Fig. 1B, right panel). The dose of puromycin selection for IFN-γ and library transfections varied from 2μg/mL to 100 μg/mL during the first month depending on cell density and viability as described in Methods. After the first month, puromycin concentration was kept at 30 μg/mL for the IFN-γ signal peptide Srt4M pool and 20 μg/mL for the signal peptide library Srt4M pool. In addition, between the first and second month, both pools were temporarily cryopreserved and stored in liquid nitrogen for 3 weeks before revival and continued selection. This selection was continued for an additional 4 months of culture. Both pools remained GFP+ during this period without an observable decrease in mean GFP intensity, demonstrating stable transfection. (Fig. 1B). Moreover, during this extended course of selection with puromycin, the width of the GFP distribution, measured by flow cytometry, narrowed, consistent with the cultures becoming more homogenous over time (Fig. 1B). We examined secretion of Srt4M from both the individual IFN-γ signal peptide stable pool as well as the signal peptide library stable pool by SDS-PAGE. We found similar Srt4M in the unpurified supernatants from both stably-transfected cell pools (Fig. 1C), and after affinity purification, confirmed positive staining for His_6_ tag by western blot (Fig. S1C).

In addition, after the 5-month selection period, we PCR amplified the signal peptide region from the stably-transfected signal peptide library Srt4M pool of cells and subjected the amplified linear fragments to next-gen nanopore sequencing (Plasmidsaurus), with results shown in Table S2. The signal peptide found at highest abundance was from Von Ebner gland protein 1 (*Rattus rattus*) followed by trefoil factor 1 (*Mus musculus*), both of which are signal peptides for proteins that are likely constitutively expressed as part of the mucosa throughout an organism’s life^51,52^.

The next most abundant signal peptide was from an antibody light chain (*Homo sapiens*) followed by fibroblast growth factor binding protein 2 (*Homo Sapiens*) and lutropin subunit beta (*Oryctolagus cuniculus*). IFN-γ signal peptide was surprisingly not found, indicating that it could have been outcompeted over time from the pool of surviving cells or the numbers of individual cells with the IFN-γ signal peptide and possibly other signal peptides from the library were too low for detection at the sequencing depth used. We conclude that multiple signal peptides, including the IFN-γ signal peptide, could enable successful secreted production of Srt4M. We anticipate these signal peptides could be useful for secreting other sortase A Δ59 variants.

Secreted expression of proteins in a mammalian host (as opposed to production in E. coli without natural glycosylation pathways) results in glycosylation if proteins have appropriate sequence motifs^53^, thus resulting in multiple products with a range of molecular weights (MW) depending on the extent of glycosylation. We observed 3 separate bands by SDS-PAGE using Ni-NTA column-purified Srt4M (Fig. 1D and S1C). The highest MW band appeared as most prevalent, the middle MW band as slightly less prevalent, and the lowest MW band was least prevalent (Fig. 1D and S1C). After adding PNGase F (New England BioLabs, NEB) to purified Srt4M, all bands collapsed to the lowest MW band (Fig. 1D). This result is consistent with the N-glycosylation of Srt4M when produced in mammalian cells, and suggests the lowest MW band represents the species without N-glycosylation.

According to prediction using NetNGlyc 1.0^53^, the highest confidence N-glycosylation site is NETR starting at position 150 of Srt4M (9/9 agreement) (Fig. S1D). Two other sites predicted with lower confidence are NESL (7/9 agreement) at position 107 and NISI starting at position 114 (5/9 agreement) (Fig. S1D). We mapped the two asparagines predicted to be glycosylated with highest confidence onto a previously reported crystal structure of sortase A and an LPETG peptide (PDB 1T2W)^54^ (Fig. S1E). Based on this structure, glycosylation at these sites seems unlikely to block enzymatic function as they do not directly overlap with LPETG peptide binding. However, the effects of glycosylation on the dynamics of a protein are hard to predict and it is difficult to make definitive conclusions without further experimentation. Nevertheless, we confirmed that the recombinant, mixed glycosylation-state Srt4M purified from Expi293 supernatants is functional by purifying it and utilizing it for the successful creation of a site-specific DAR1 antibody-siRNA conjugate as presented later in the text.

### Stable production of antibodies with heavy-chain tags

To enable production of a complete antibody molecule comprised of heavy and light chains, we used a tricistronic cassette containing an immediate early CMV enhancer/promoter and an additional WT EMCV IRES (as described in Ho et al. in CHO cells^26,27^). The cassette is still linked to the attenuated EGFP-P2A-PuroR selection cistron as discussed above. Within this tricistronic cassette, the antibody light chain is the first cistron translated by cap-dependent translation. The second cistron is the antibody heavy chain, translated by WT EMCV IRES-mediated translation. Finally, EGFP-P2A-PuroR is the third cistron, translated by attenuated IRES-mediated translation. In addition to this tricistronic cassette, the complete expression vector, as shown in Fig. 2A, contains the same S/MAR and IR elements already described earlier in the text for episomal maintenance in Expi293 cells (Fig. 2A). An expanded cloning protocol is described in Methods.

**Figure 2.**
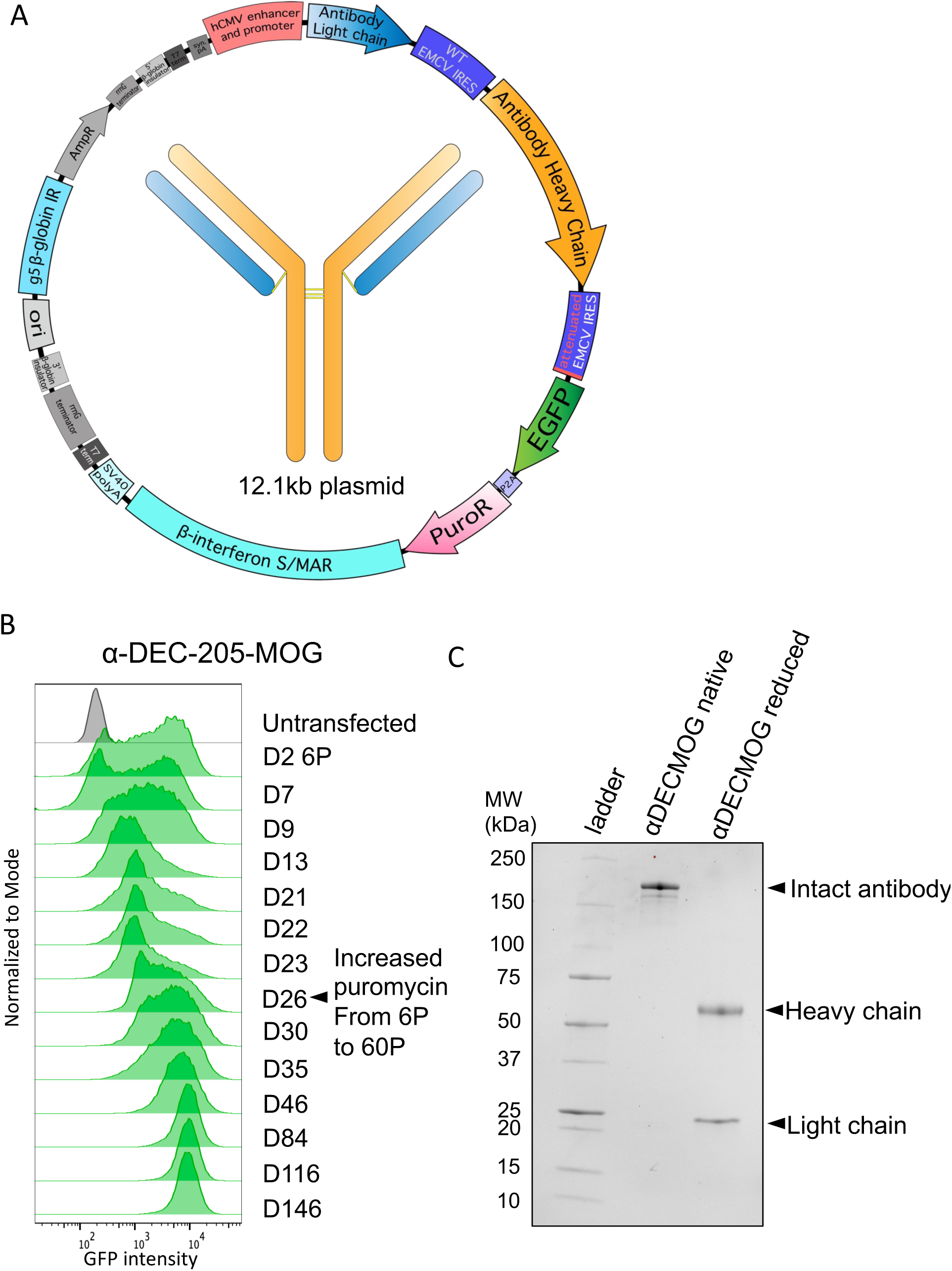
Generation of a stable cell pool expressing αDEC-205-MOG. **A.** Tricistronic plasmid map for antibody expression. Elements include, as indicated: hCMV promoter and enhancer, antibody light chain, WT EMCV IRES, antibody heavy chain, attenuated EMCV IRES, EGFP, P2A self-cleaving peptide, PuroR, β-interferon S/MAR, SV40 pA, G5 β-globin IR **B.** Cells were transfected with a tricistronic plasmid as in A that included heavy and light chains of αDEC-205-MOG. Histograms show GFP expression intensity over a time course ranging from 2 days to 146 days after transfection. Cells were initially selected at 6 μg/mL puromycin (6P), which was later increased to 60 µg/mL (60P) on D26 after transfection as indicated. **C.** SDS-PAGE gel showing protein G purified-αDEC-205-MOG with and without disulfide bond reduction performed before loading (reduced and native, respectively)

To examine the detailed kinetics of the process of stable pool generation using the vector with a tricistronic cassette, we expressed a chimeric antibody specific for mouse DEC-205 which contains myelin oligodendrocyte glycoprotein (MOG)_35-55_ peptide fused to the heavy chain C-terminus (αDEC-205-MOG) with a linker as previously described^55^. As seen in Fig. 2B, an initial GFP^+^ pool was generated by selection with 6 µg/ml puromycin in approximately 23-26 days. To enable an accumulation of higher producing clones, the puromycin dose was then increased to 60 µg/ml. This resulted in a dramatic increase in mean GFP intensity that prevailed long-term and was stably maintained for at least 5 months of continuous culture with selection (Fig. 2B). After completing the selection, we confirmed the production of the complete antibody that included both light and heavy chains by SDS-PAGE after protein G purification from supernatants (Fig. 2C). Overall, we achieved successful stable expression of a C-terminally tagged antibody with the episomally replicating vector using a tricistronic cassette with attenuated expression of EGFP-P2A-PuroR.

### Utilizing inteins for production of a bispecific antibody-like molecule

For specific pairing of an antibody heavy chain with a protein-Fc fusion chain, we utilized complementary electrostatic steering mutations previously described by Wang et al. and established to result in 100% correct heavy chain pairing^56^. This approach involves point mutations E356K and D399K in one heavy chain CH3 domain and the complementary K439D and K409E mutations in the other heavy chain CH3 domain. These complementary mutations break the natural heavy chain symmetry and allow only charge-complementary pairing to occur^56^. We further combined this complementary charge approach based on specific mutations in different heavy chains with the SiMPl split intein-split puromycin system^39^ to obtain a complete expression system for an immunoglobulin molecule with two different heavy chains, as described in further detail below.

Split inteins can be used to reconstitute, by autocatalytic formation of a peptide bond, a protein that has been designed as two separate, dysfunctional parts^39,57^. Specifically, two dysfunctional fragments of PuroR protein, expressed under two separate cassettes^39^, can be re-formed using split inteins^39^. We split the EGFP-P2A-PuroR cistron, originally present in a single expression cassette described in Figs. 1A and 2A into two cistrons included in two separate expression cassettes. Specifically, we included EGFP-P2A-N-terminal-PuroR-N-terminal-intein in one cassette and C-terminal-intein-C-terminal-PuroR in another cassette (Fig. 3A). In these two new expression cassettes, both intein-PuroR fragments are still expressed by attenuated IRES-mediated translation following the expression of the corresponding mutated heavy chain (Fig. 3A and B). Therefore, for both intein-PuroR fragments to be expressed, to associate, and to splice together the functional PuroR, the expression of both complementary heavy chains is required. The individual expression cassettes can either be co-transfected as part of two separate plasmids (Fig. S2A) or transfected together as part of a single plasmid with two promoters (Fig. S2B), thus completing the episomal vector design. Expanded cloning details are included in Methods. To demonstrate the utility of this system, we designed an αDEC-205*-*programmed cell death protein 1 (PD-1)*-*Fc bispecific construct (Fig. 3B). The αDEC-205 light chain and heavy chain (with K409E and K439D mutations) were expressed from a tricistronic cassette with the C-terminal intein C-terminal PuroR vector (Fig. 3B right panel) and the mouse PD-1-Fc chain (with E356K and D399K mutations) was expressed from a bicistronic cassette with the N-terminal PuroR N-terminal intein vector (Fig. 3B left panel). We used a linker design (IgG3C-) similar to that pioneered by Bournazos et al.^58^ for extended flexibility between both immunoglobulin arms, with a cartoon backbone representation of an atomic model shown in Fig. 3C. We co-transfected both plasmids and achieved a stable pool in about 1.5-2 months. The PD-1-Fc chain also included a His_6_-tag that allowed the entire αDEC-205*-*PD-1-Fc molecule to be purified by Ni-NTA column. We purified αDEC-205*-*PD1*-*Fc using a Ni-NTA column and examined it by SDS-PAGE in native (non-reduced) and reduced forms with or without an additional deglycosylation step. We observed bands corresponding to the expected MW of 53.2 kDa for αDEC-205 heavy chain and 45.0 kDa for PD-1-Fc in the sample that was deglycosylated and run under reducing conditions. In addition, we observed a band consistent with an expected MW of 121.6 kDa for the entire αDEC-205*-*PD1-Fc molecule after its deglycosylation without disulfide reduction (native) (Fig. 3D). In contrast, without the deglycosylation step, both native and reduced forms ran at higher than predicted MWs, consistent with the glycosylation of both heavy chains (Fig. 3D). Overall, we conclude that intein-mediated selection coupled with complementary electrostatic steering mutations allows successful, stable production of a bispecific antibody-like molecule.

**Figure 3.**
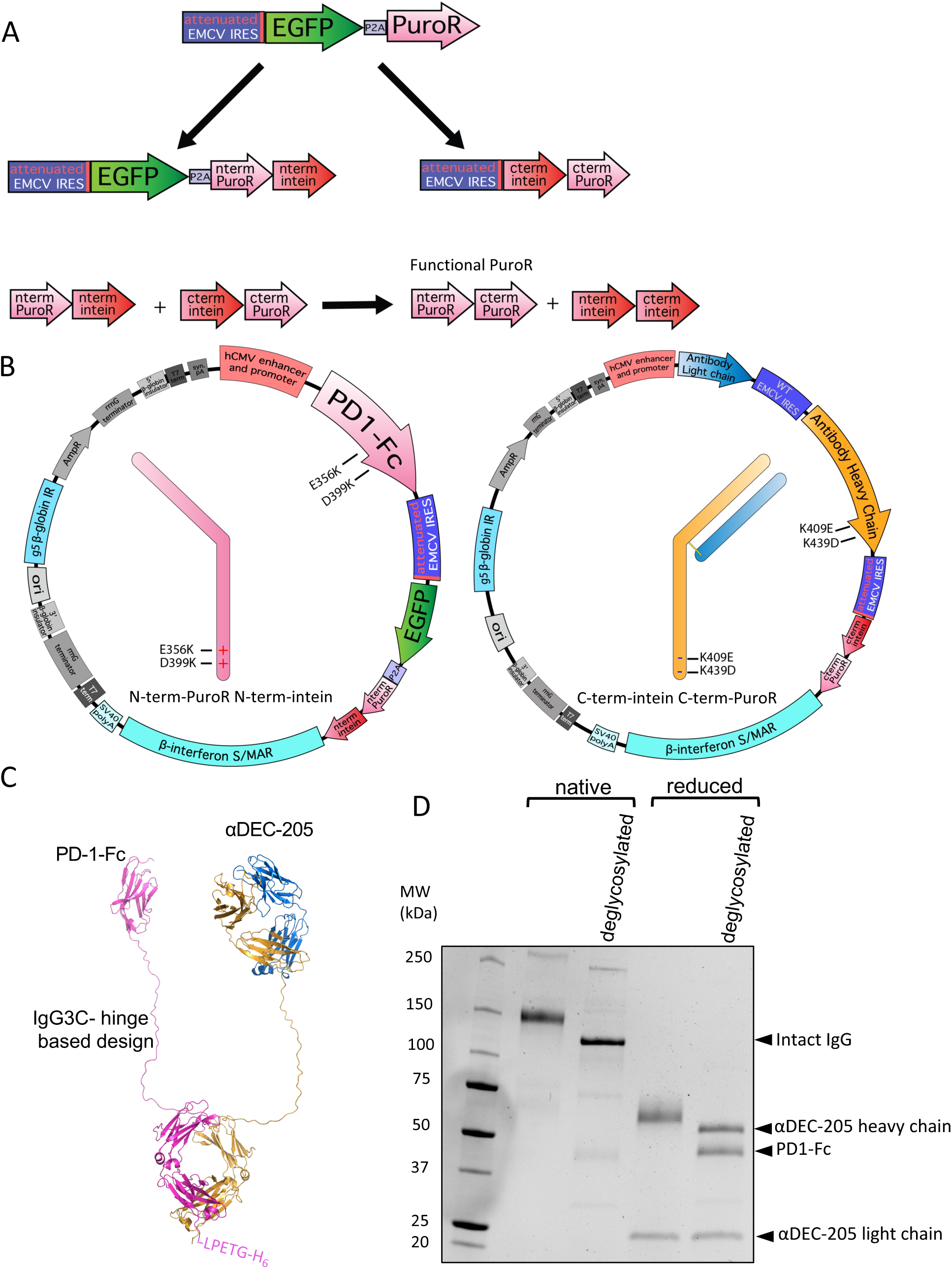
Split intein-mediated selection to produce a bispecific αDEC-205-PD-1-Fc molecule. **A**. Diagram showing intein-mediated splitting of puromycin based on the SiMPl system^39^. The EGFP-P2A-PuroR cistron is split into two cistrons: EGFP-P2A-N-terminal-PuroR-N-terminal-intein cistron and C-terminal-intein-C-terminal-PuroR cistron, both translated via an attenuated IRES as shown. When expressed together, both intein-PuroR fragments reform functional PuroR, as indicated. **B**. Plasmid maps showing the linked intein-PuroR fragments with specific heavy chain mutations. E356K and D399K within PD-1 Fc are linked to the N-terminal-PuroR N-terminal-intein cistron (left). K409E and K439D within the αDEC-205 heavy chain are linked to the C-terminal-intein C-terminal-PuroR cistron (right). **C**. Cartoon backbone representation diagram of an atomic model of αDEC-205-PD-1-Fc bispecific antibody-like molecule. The structure was first generated with Alphafold multimer 2.3.2^70,71^. Rotations of each arm were then made in PyMOL (Schrödinger) for display. Disulfide bonds between the two heavy chains were added in BioLuminate (Schrödinger) and the molecule was minimized. Image generated using PyMOL (Schrödinger) **D**. SDS-PAGE gel showing αDEC-205-PD-1-Fc after nickel column purification with and without disulfide bond reduction before loading (reduced and native, respectively) and additionally, with and without deglycosylation mix II (NEB) treatment (deglycosylated as indicated). Arrows on side indicate the bands of expected MW as described in the text representing the full bispecific molecule and individual heavy chains when deglycosylated.

### Split inteins for stable production of defined multi-tag antibodies

Intein-mediated selection coupled with complementary electrostatic steering mutations can also be applied to produce antibodies with distinct C-terminal modifications in their heavy chains. Following the strategy discussed in Fig. 3 above, we designed αDEC-205 antibodies where one heavy chain (with K409E and K439D mutations) included an antigenic fragment from ovalbumin (OVA_323-339_) with a linker as previously published^59^ while the other heavy chain (with E356K and D399K mutations) included the same linker followed by a sortase tag and His_6_ tag (LPETGH_6_). Transfecting these two cassettes linked to attenuated IRES-mediated expression of split intein-split puromycin fragments allows the creation of antibodies with an antigen and a single sortase attachment site (Fig. 4A). We also separately generated stable Expi293 pools expressing heavy chain sequences without complementary mutations that have either OVA_323-339_ or LPETGH_6_ on both chains with the same linker. A depiction of these antibodies is shown in Fig. 4B. Because the heavy chain with an LPETGH_6_ tag has a MW 1.5 kDa lighter than the MW of the heavy chain with OVA_323-339_, all three produced antibodies could be readily distinguished from each other by SDS-PAGE when disulfides are reduced (Fig. 4C). Two bands of different MW but equal intensity are seen in the αDEC-205-ΟVA-LPETGH_6_ sample, as expected for this molecule that contains two different heavy chains. In contrast, and as expected, only a single corresponding heavy chain band is present in the case of αDEC-205-ΟVA or αDEC-205-LPETGH_6_ antibodies (Fig. 4C). Overall, we conclude that by attenuated IRES-linked split intein-mediated selection, we successfully achieved stable production of an antibody molecule that contains different C-terminally modified heavy chains, including a single sortase tag and a single antigen molecule.

**Figure 4.**
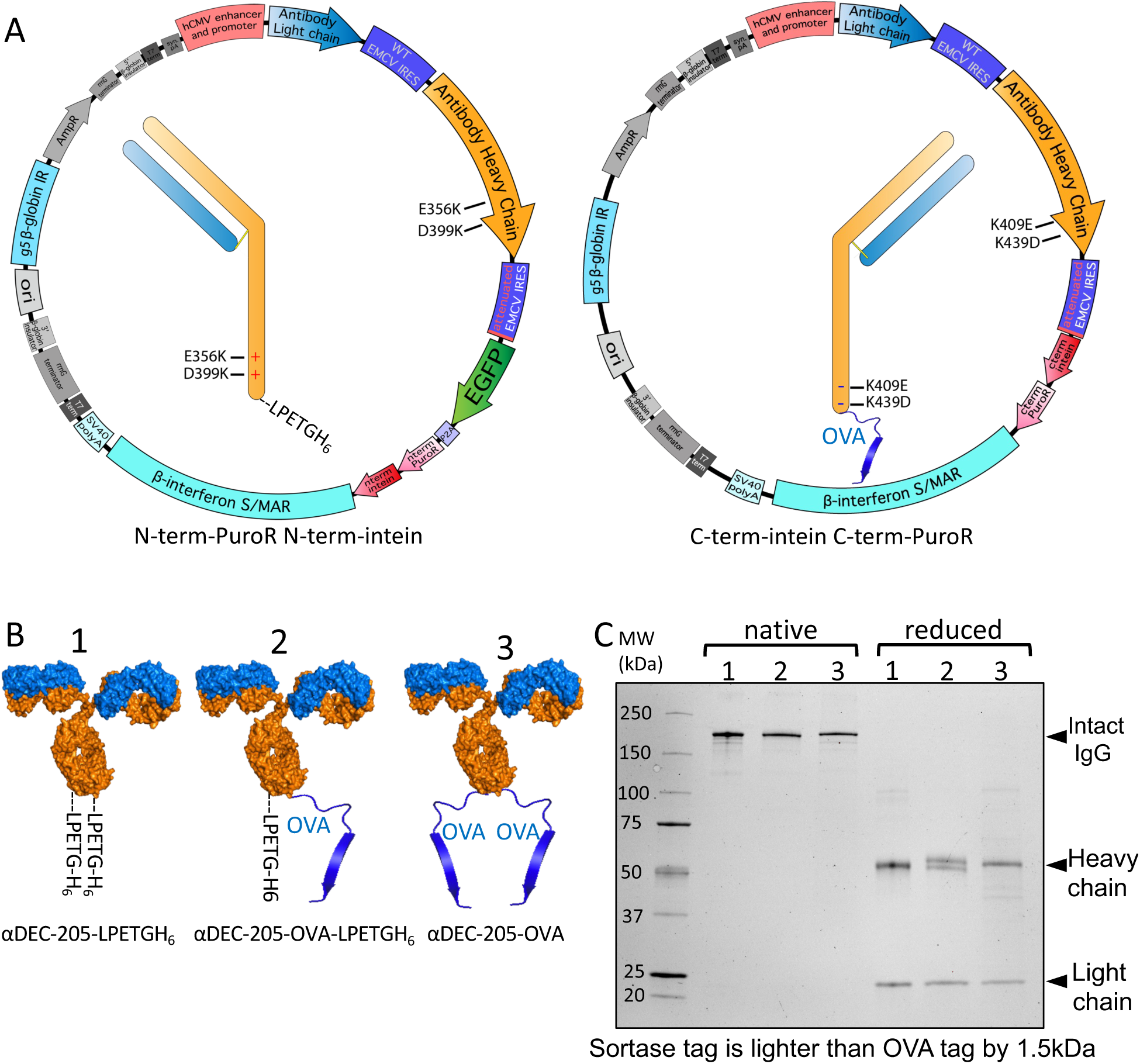
Split intein-mediated selection to produce antibodies with two different C-terminal tags. **A.** Plasmid map showing coupling of LPETGH_6_ tag on one heavy chain (with E356K and D399K mutations) with EGFP-P2A-N-terminal-PuroR-N-terminal intein (left) and coupling of OVA_323-339_ antigen on another heavy chain (with K409E and K439D mutations) with C-terminal intein-Cterminal-PuroR (right). **B.** Diagram of antibodies as numbered: 1-αDEC-205 antibody with a sortase tag and His_6_ tag (LPETGH_6_) on both chains. 2 - αDEC-205 antibody with LPETGH_6_ on one chain and OVA_323-339_ antigen on the other chain. 3- αDEC-205 antibody with OVA_323-339_ antigen on both chains. **C.** SDS-PAGE gel showing native and reduced antibodies from B as indicated by the numbers on top of the lanes. Native antibodies are loaded without disulfide bond reduction, while reduced samples have had disulfide bonds reduced before loading.

### Site-specific DAR1 siRNA-antibody conjugate

To show the applicability of an antibody produced with a single sortase tag, we created a site-specific DAR1 siRNA-antibody conjugate. Based on the approach we presented in Fig. 4, we generated an antibody that contained a single sortase tag (LPETGH_6_) directly on the C-terminus of one heavy chain, without any linkers. Next, we made use of the recombinant Srt4M that we purified from Expi293 supernatants (Fig. 1). An outline of the process of sortagging a GGG-PEG_4_-DBCO molecule to an antibody with a single LPETGH_6_ tag is shown in Fig. 5A and additional details are included in Methods. Approximately 56 μg of Srt4M was added to 150 μg of antibody with excess GGG-PEG_4_-DBCO and 5 mM CaCl_2_ in a total volume of approximately 200 μL. After 15 minutes, EDTA was added to quench the reaction and the antibody, Srt4M, and excess GGG-PEG_4_-DBCO were separated by size exclusion chromatography (SEC) (Fig. 5B). A thiol-modified siRNA was conjugated to an azide-containing molecule using Azido-PEG_3_-Maleimide according to the manufacturer’s protocol (Fig. 5C). The modified siRNA was also separated from free Azido-PEG_3_-Maleimide by SEC (Fig. 5D). The purified DBCO-modified antibody and the purified azide-modified siRNA were then mixed overnight at 4°C (Fig. 5E) in a click chemistry reaction^40^ and resolved by SDS-PAGE the following day (Fig. 5F). A clear shift in MW (consistent with the ∼14 kDa MW siRNA) was seen in the sample conjugated with siRNA without disulfide bond reduction. Analysis under reducing conditions further revealed the expected presence of both the siRNA-conjugated and unconjugated heavy chains (Fig. 5F). This is an expected result as only one of the two heavy chains has a sortase tag and therefore, the ability to be conjugated. The similar relative intensities of the conjugated and unconjugated heavy chain bands indicate a high degree of conjugation of the DBCO-sortagged antibody. Additionally, we visualized the siRNA-conjugated antibody by resolving it on another gel that was imaged after incubation with ethidium bromide (EtBr), which intercalates double stranded nucleotides like siRNA (Fig. 5G). Despite a small amount of visible non-specific protein staining by EtBr, we observed a striking difference in band intensity corresponding to the siRNA-conjugated antibody and heavy chain, further confirming the presence of nucleic acid conjugation (Fig. 5G). Overall, we conclude that we successfully created a DAR1 siRNA-antibody conjugate by utilizing the key reagents that were all produced in Expi293 cells using our novel expression vectors and strategies.

**Figure 5.**
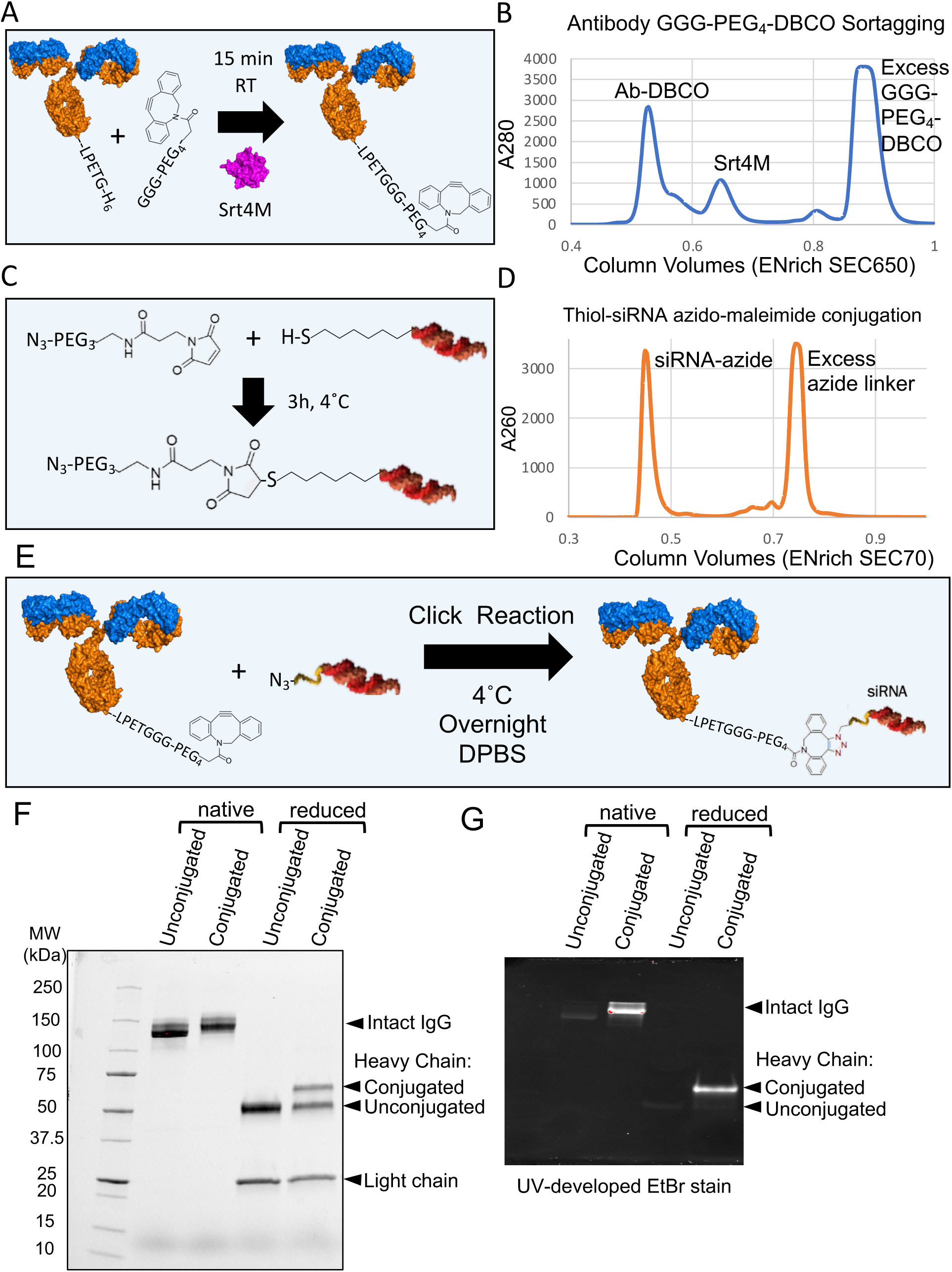
Creation of DAR1 antibody-siRNA conjugate by sortagging and click chemistry. **A.** Diagram of DBCO-PEG_4_-GGG mediated sortagging of antibody with a single sortase tag. GGG-PEG_4_-DBCO is sortagged onto the antibody using Srt4M as shown. **B.** A280 chromatogram showing separation between antibody-DBCO conjugate, Srt4M, and free GGG-PEG_4_-DBCO linker using an ENrich SEC 650 column (BioRad). **C.** Diagram of thiol-modified siRNA conjugation to azido-PEG_3_-maleimide. The maleimide group reacts with the free thiol on reduced siRNA to form the azide-modified siRNA conjugate as shown. **D.** A260 chromatogram showing separation of siRNA-azide conjugate from excess azide-maleimide linker using an ENrich SEC 70 column (BioRad). **E.** Diagram outlining formation of antibody-siRNA site-specific DAR1 conjugate using DBCO-azide-based click chemistry as shown. **F.** 4-20% gradient SDS-PAGE gel of antibody-siRNA conjugate. Unconjugated and conjugated antibody with and without disulfide bond reduction before loading (reduced and native respectively) are shown as indicated. **G.** 7.5% SDS-PAGE gel of antibody-siRNA conjugate after gel staining with EtBr. Same sample loading order as in F. Unconjugated and conjugated antibody with and without disulfide bond reduction before loading (reduced and native respectively) are shown as indicated.

## DISCUSSION

We present a comprehensive plasmid system useful for stable long-term production of recombinant proteins, standard antibodies, modified antibodies, and bispecific antibody-like molecules. By combining episomal retention with split selection of two separate protein expression cassettes, our approach represents a significant advancement for stable production of complex molecules in Expi293 cells. We also demonstrated a novel approach to identify signal peptides that allow secretion of Srt4M in mammalian cells. Overall, our novel expression systems allowed us to produce recombinant proteins that we used to prepare a site-specific DAR1 antibody-siRNA conjugate.

We started with the general tricistronic cassette design as described by Ho et al. over a decade ago^26^. In this system, the expression of both antibody chains is linked to a single selectable marker expressed from an attenuated IRES. Critically, this tricistronic expression system allows greater protein productivity and increases the likelihood of stable production of both chains over time^26–28^. However, this tricistronic approach based on a single antibiotic is limited to stable expression of only one specific light chain and heavy chain. In this work we present a major breakthrough for the stable production of more complex antibody molecules using two different tricistronic cassettes in combination with a split selectable marker. Specifically, each individual tricistronic cassette is linked to the expression of a split-intein-split-PuroR-fusion protein, which on its own is not a functional selectable marker. It is only when two tricistronic cassettes are expressed together that both intein-PuroR fragments are expressed and can re-form the functional selectable marker. Therefore, this system permits the expression of up to 4 different protein chains linked to selection by a single selectable marker, unlocking a wide range of possibilities for the stable production of complex molecules, the full scope of which we have only just begun to explore in this paper.

We demonstrated the utility of paired tricistronic cassettes for achieving stable production of antibodies with two different C-terminally modified heavy chains. We also showed an example of a tricistronic cassette paired with a bicistronic cassette encoding a protein-Fc fusion to stably produce an antibody-protein-Fc bispecific molecule. We anticipate that paired tricistronic cassettes should enable the production of classical bispecific antibodies, with two different light chain-heavy chain pairs. For example, highly specific approaches such as CrossMab^CH1-CL^ or similar methods that produce no incorrect light chain-heavy chain pairing^60–62^ could work well in a tricistronic format where light chains are expressed slightly in excess to heavy chains^63^.

While paired tricistronic cassettes have a clear advantage in enforcing the expression of 4 different protein chains by a single antibiotic, they are not strictly required to take advantage of attenuated IRES-mediated split intein selection. We expect that the vectors we generated would enable expression of a single light chain and heavy chain from separate promoters as bicistronic cassettes linked to split-selection with a single antibiotic. However, to adapt this approach to bispecific antibody expression, with each chain expressed under an independent promoter, would require either further splitting a single selection marker into 4 fragments or using two different split selection markers, such as split-puromycin and split-hygromycin, previously generated by Palanisamy et al^39^.

Biotherapeutic manufacturing requires yield optimization by a variety of methods including deriving and screening clonal lines and production by high density (15-20 x 10^6^ cells/mL) fed-batch culture. In this work, we focused on the development of methods for reliable production of proteins at the preclinical stage of development. Our approach ensures that all protein chains are expressed upon long-term selection to enable the stable production of complex molecules. In addition, the inclusion of EGFP together with PuroR within the selection cistron provides the crucial ability to regularly monitor protein expression during the selection process. Furthermore, this design allows live cells to be selected by fluorescence activated cell sorting. Therefore, our approach for the generation of stably transfected pools of cells is likely to be useful as a starting point for preclinical protein production and may also find use for production of biotherapeutics.

Our novel expression systems enabled us to make products that allowed the creation of a DAR1 site-specific antibody-siRNA conjugate. We anticipate a very similar strategy could be used to conjugate other nucleic acids, such as Toll-like receptor-activating adjuvants. The antibody-based delivery of various antigens to dendritic cells has now been well-established and broadly applied for modulating specific T cell responses^36,64^. The current generation of dendritic-cell targeted antigen therapeutics in human cancer trials utilize a combination of many components including non-specifically delivered adjuvants such as Poly-ICLC and resiquimod^37,38^. We anticipate the approaches presented in this work could ultimately allow the delivery of nucleic acid-based adjuvants directly conjugated to engineered antibodies containing specific antigens. We also speculate that the use of other bond-forming enzymes^49,65–68^, in addition to Srt4M, could make it possible to conjugate multiple antigens^69^ along with an adjuvant to antibodies in a site-specific manner. These approaches could enable the next-generation of dendritic cell-targeted antigen and neoantigen cancer therapeutics.

In closing, we present a robust and comprehensive expression system in Expi293 cells based on episomal replication with protein expression directly linked to selection. We use this system to reliably produce recombinant proteins and engineered antibodies in stably-transfected cells. We also demonstrated the applicability of the produced reagents by making a site-specific antibody-nucleic acid conjugate. Overall, these novel approaches and methods are likely to be of a very high interest for research, preclinical, and industrial applications.

## ACKNOWLEDGMENTS

The authors would like to thank Joy Eslick and Jackie Spencer (Saint Louis University Flow Cytometry Core) for expert help with flow cytometry. We thank the Saint Louis University Institute for Drug and Biotherapeutic Innovation for support with chromatography. The authors also thank John Walker, Saint Louis University, for assistance with the Azido-PEG_3_-Maleimide protocol, Ziva Misulovin, Saint Louis University, for providing cloning reagents, and James Brien, Saint Louis University, for helpful discussions. This work was supported in part by grants from National Institute of Allergy and Infectious Diseases of the National Institutes of Health (R01AI113903), National Multiple Sclerosis Society (RG-1902-33632), and National Multiple Sclerosis Society (RFA-2104-37543), all to DH.

## AUTHOR CONTRIBUTIONS

I.F. designed, cloned, produced, purified, and characterized all molecules presented here and wrote the manuscript. D.H. oversaw the project and wrote the manuscript. Both authors contributed to the interpretation of the results and direction of work.

## MATERIALS AND METHODS

All plasmid vector backbones will be available from Addgene upon publication. The tricistronic EGFP-2A-PuroR vector backbone is available at Addgene ID 208377. The tricistronic N-terminal-Puro-N-terminal-intein vector backbone is available at Addgene ID 208378. The tricistronic C-terminal-Puro-C-terminal-intein vector backbone is available at Addgene ID 208379.

### Vector Construction

The original backbone CMV expression vector was obtained from Twist Bioscience. Whenever possible, sequences were ordered from Twist Bioscience or Integrated DNA Technologies. All sequences were codon optimized for expression in human cells using the built-in codon optimizer tools from each company. Further, all ordered sequences excluded restriction enzymes NotI, EcoRI, BstXI, NheI, XmaI, SalI, XbaI, FseI, AscI, SbfI, and MluI except at their designated positions, to maintain flexibility of the plasmids. For sequences that could not be synthesized due to repeated nucleotides or other internal complexities such as EMCV IRES, β-globin IR, and beta-interferon S/MAR, they were PCR amplified using Accuprime Pfx (Invitrogen) as follows: EMCV IRES was amplified from plasmid pEP4 E02S EN2L^72^, which was a gift from James Thomson (Addgene plasmid #20922; http://n2t.net/addgene:20922; RRID:Addgene_80391). β-globin IR was amplified from BAC clone RP11-1205H24 obtained from BACPAC Genomics Inc. β-interferon S/MAR was amplified from plasmid pBMN(CMV-copGFP-Puro-SMAR)^73^, which was a gift from Magnus Essand (Addgene plasmid #80391; http://n2t.net/addgene:80391; RRID:Addgene_80391). The split puromycin intein sequences were synthesized by Twist Bioscience based on the sequences in Addgene plasmids 134319 and 134319^39^.

### Cloning of Single Proteins (e.g. Srt4M) Plasmids

The EGFP-P2A-PuroR vector backbone contains unique restriction sites for cloning by restriction enzyme digestion and ligation (Fig. S1A). For cloning of single proteins such as Srt4M, we ordered a DNA fragment encoding Srt4M with upstream NotI (underlined) and Kozak sequence (italics) before the start codon “<u>gcggccgc</u>*gccgccacc*atg” with an NheI site added after the stop codon. The insert and the EGFP-P2A-PuroR vector backbone (Fig. S1A) were digested with NotI and NheI, appropriate fragments were gel purified, ligated together for 30 min at room temperature according to standard NEB ligation protocol, and transformed into DH5α or NEB Stable.

### Cloning of Antibody Plasmids

Antibodies vectors are cloned by a 4-part ligation including: light chain, WT IRES, heavy chain, and the vector backbone. The light chain sequence was ordered with upstream NotI (underlined) and Kozak (italics) before the start codon “<u>gcggccgc</u>*gccgccacc*atg” with an EcoRI site added after the stop codon. The beginning of the heavy chain sequence was ordered in frame with a BstXI site as ccacaacc<u>atg</u>g (underlined atg is the heavy chain signal peptide start codon) with an NheI site added after the stop codon. The WT IRES fragment (between EcoRI and BstXI sites) can be obtained by digesting any of vector backbones. The desired vector backbone, either EGFP-P2A-PuroR (Fig. S1A), N-terminal intein or C-terminal intein plasmid (Fig. S2A), is then digested with NotI and NheI. The light and heavy chains are digested with corresponding enzymes as described above. All 4 gel purified fragments (light chain, heavy chain, IRES, vector backbone) were ligated together at room temperature for 2 hours according to standard NEB protocols and then transformed into DH5α or NEB Stable.

### Single Plasmid for Bispecific Production

To obtain a single vector, we digested the N-terminal intein plasmid (Fig. S2A – left panel) with MluI and SbfI to obtain the vector backbone after gel purification. We also digested the C-terminal intein plasmid (Fig. S2A – right panel) with MluI and SbfI to excise the expression cassette that was also gel purified. We ligated together for 30 min at room temperature using standard NEB protocols and transformed into NEB Stable (30°C). This allows the efficient formation of a combined vector (Fig. S2B) containing both tricistronic cassettes linked to N- and C-terminal intein-puromycin cistrons, with each cassette containing an S/MAR and the entire plasmid containing a single IR. This combined vector loses many unique restriction sites and therefore cannot easily be used for further cloning. It is instead used directly for transfection as a single plasmid containing both the N-terminal tricistronic cassette and the C-terminal tricistronic cassette.

### Cells, Transfection, and Selection

Expi293F cells (A14527 Gibco) were grown in a humidified 8% CO2 incubator. For culture volumes 20 mL or more, cells were shaken at 125 rpm. For smaller culture volumes of 10 mL or less, cells were shaken at an angle at speeds between 125-145 rpm in 50 mL bioreactor tubes.

Cell viability and density were measured using a Countess II FL Automated Cell Counter (Invitrogen). DNA for transfection was purified using EndoFree Plasmid Maxi Kit (Qiagen). For single plasmid transfections, we used 75 μg DNA per 60 x 10^6^ cells at a density of 20 x 10^6^ cells/mL with a polyethylenimine (PEI):DNA ratio of 3:1 and 0.1% Pluronic F-68 in Expi293 media for 3 hours as described in Fang et al^74^. As also described in Fang et al., more DNA can be used but PEI should be kept at 225 μg per 60 x 10^6^ cells. For co-transfection of two plasmids, we used 375 μg of DNA per 200 x 10^6^ cells, split between the two plasmids (187.5 μg DNA/200 x 10^6^ cells per bispecific plasmid) while keeping PEI at 750 μg per 200 x 10^6^ cells. After transfection, cells were diluted to approximately 1 x 10^6^ cells/mL, typically in an equal mixture of Expi293 and EX-CELL 293 (14571C Sigma). Expi293 media can be used for the entire process; however, to reduce media expenditures, we typically cultured cells in a mixture of both media during selection and eventually adapted them to growing only in EX-CELL 293 media (after 3-4 weeks). Puromycin (Invivogen) was typically added at 2-6 μg/mL 24-48 hours after transfection. During the first month after transfection, the puromycin concentration was sometimes kept at 2-6 μg/mL and was sometimes adjusted based on cell viability and density.

Typically 2μg/mL is sufficient to generate stable pools, but we tended to increase selection pressure as cells continued to expand. When cell density was above 1 x 10^6^ cells/mL, higher concentrations of puromycin were temporarily used (typically 20-60 μg/mL) compared to when cell density dropped below 1 x 10^6^/mL. When cell density dropped below 3 x 10^5^ cells/mL, 0-2 μg/mL puromycin was temporarily used (from 12hr to a few days) until the cells recovered and then higher dose selection is resumed. If selection is removed for too long, low dose resistance to puromycin can emerge. This can be resolved either by increasing the dose of selection or sorting for GFP^+^ cells. We define a cell pool as stable at a particular dose of puromycin when after seeding at 3-5 x 10^5^ cells/mL into fresh selective media, cells double approximately every 24 hours while maintaining >95% viability without changes in mean GFP intensity. Selection with puromycin is maintained during regular passaging, however, before final production, selection is removed for at least 2 passages to ensure the least stress on the cells.

### Antibody Expression and Protein Purification

For protein expression, cells were seeded at a density of 1.5 x 10^6^ cells/mL, typically in EX-CELL 293 media (though Expi293 media supports higher cell densities/yields) and valproic acid was added to the culture at a concentration of 3.5mM as described in Fang et al^74^. After about 5 days (or when viability drops below 50%), supernatants were spun down successively at 250g, 1600g, and 4600g for 30 minutes followed by sterile filtering with a 0.22 µm filter. Filtered supernatants were applied directly onto a column. Alternatively, large volumes can be further concentrated and dialyzed into DPBS using a Vivaflow 50R crossflow cassette (Sartorius) before column purification. Antibodies were purified by gravity column using Protein G Sepharose 4 Fast Flow Resin (Cytiva) and proteins containing His tags were purified on Ni-NTA Agarose (R90101 Invitrogen).

### Sortase Signal Peptide Library Construction

The entire secreted mammalian proteome was downloaded from UniProt, and proteins including signal peptides were sorted based on the first 2 amino acids after the predicted signal peptide cleavage site (according to UniProt). Peptides from proteins with QA sequence (matching the QA of Srt4M Δ59) were initially selected and further curated manually to obtain 17 signal peptides that were chosen to be part of a signal peptide library (Table S1). Three DNA fragments were ordered from Twist Bioscience that each contained sequences of 5-6 signal peptides along with in frame restriction enzyme sites (NotI and BlpI). Each fragment was digested with NotI and BlpI and the mixture of individual DNA fragments encoding signal peptides were ligated into the plasmid with Srt4M described above and transformed into DH5α. For each of the three ligations (with no visible control ligation colonies), around 40 colonies were selected, grown together in liquid culture, purified by EndoFree Maxiprep (Qiagen), then transfected into Expi293 cells. Puromycin (Invivogen) was added after the first 48 hours. When cell density was above 1 x 10^6^ cells/mL, higher concentrations of puromycin were used (60-100 μg/mL) compared to when cell density dropped below 1 x 10^6^/mL. When cell density dropped below 3 x 10^5^ cells/mL, 0-2 μg/mL was temporarily used until the cells recovered. After the first month, 20 μg/mL puromycin was used. For deglycosylation experiments, PNGase F (P0710S NEB) was used. For western blot to detect His_6_ tag, we used HRP anti-His tag antibody (BioLegend 652503).

### Signal Peptide Library PCR Amplification and Sequencing

Cells were incubated at 52°C in Proteinase K-containing detergent buffer. PCR primers (F primer 5’ agatcagatctttgtcgatcctacca 3’ and R primer 5’tcatctcccggggttgtggc 3’) were used to amplify the entire first cistron and IRES using Accuprime Pfx (Invitrogen) for 30 cycles. The amplified fragments corresponding to the expected MW were gel purified and sent to Plasmidsaurus for standard Oxford Nanopore sequencing of linear fragments. Only complete reads including the entire signal peptide to the beginning of Sortase A without errors were considered in the analysis.

### Antibody and Protein-Fc Constructs

αDEC-205 constructs were previously described in Hawiger et al.^34^ αDEC-205-OVA and MOG sequences at C-terminus with linkers are as previously described ^55,59^. αDEC-205-OVA-LPETGH_6_ included LPETGH_6_ in place of MOG or OVA antigens. αDEC-205 with a single sortase tag (Fig 5) included LPETGH_6_ directly fused to the C-terminus of one heavy chain without any linker. For bispecific αDEC-205-mouse PD-1-Fc, a mutant IgG3 linker was used (IgG3C-)^58^ with two repeats (<u>ELK</u>SPRSPEPKSSDTPPPSPRSPEPKSSDTPPPCPRCPAPEL).

An Ig light chain signal peptide (MAWTPLLLPFLTLCIGSVVS) was used to express mouse PD-1 residues 25-150 (C83S) with (ERIL<u>ELK)</u> as the overlap to the IgG3 hinge. For deglycosylation experiments, deglycosylation mix II (P6044S NEB) was obtained from New England Biolabs (NEB). Some deglycosylation enzymes can be seen on SDS-PAGE gels when added.

### siRNA

The siRNA was purchased from TriLink Biotechnologies with HPLC purification. The sequences were based on Katakowski et al.^75^ but with additional modifications as described below: The antisense siRNA sequence that was ordered was as follows: 5’ [Phos(H)] A(ps)[fA](ps)G[fU]C[fG]U[fA]G[fA]G[fU]C[fC]A[fG]U[fU]G(ps)U(ps)U 3’ where all bases were 2’ O-Methyl modified unless indicated in brackets, [fX] indicates 2’ Fluoro Base,[Phos(H)] indicates 5’ Phosphate,(ps) indicates phosphorothioate bond. The sense siRNA strand was as follows: 5’C(ps)A(ps)[fA]C[fU]G[fG]A[fC]U[fC]U[fA]C[fG]A[fC]UU[SSC6] 3’ where all bases were 2’ O-Methyl modified by default unless indicated in brackets, [fX] indicates 2’ Fluoro Base, (ps) indicates phosphorothioate bond, [SSC6] indicates 3’ C6 Disulfide Linker. Individual siRNA strands were dissolved in H_2_O at approximately 6 μg/μL. They were then mixed together equally and annealed by heating to 90°C for 1 min, then placed in a 70°C water bath and allowed to cool gradually over an hour with the water bath turned off. After annealing, siRNA was treated with excess TCEP (to reduce the C6 disulfide linker) and then buffer exchanged back into water using a Pierce Protein concentrator with 3K MWCO.

### Antibody-siRNA Conjugation

Approximately 56 μg of Srt4M (freshly buffer exchanged into DPBS to remove TCEP that is contained in the Srt4M stock as described in Li et al.^29^) was added to 150 μg of antibody with excess GGG-PEG_4_-DBCO and 5 mM CaCl_2_ in a total volume of approximately 200 μL. After 15 minutes, 20 mM EDTA was added to quench the reaction and the antibody, Srt4M, and excess GGG-PEG_4_-DBCO were separated by SEC using an ENrich SEC 650 column (BioRad) (Fig. 5B) on a BioRad NGC Quest 10 Plus. Azido-PEG_3_-Maleimide was prepared according to the manufacturer’s protocol under Argon gas in dry DMSO. Azido-PEG_3_-Maleimide (in DMSO) was added in significant excess to a thiol-modified siRNA (as prepared in the previous siRNA methods section) and left at 4°C for 3 hours. The modified siRNA was then separated from free Azido-PEG3-Maleimide by SEC using an ENrich SEC 70 column (BioRad) on a BioRad NGC Quest 10 Plus. Fractions containing the DBCO-sortagged antibody and azide-modified siRNA were concentrated using Pierce or Amicon concentrator columns (3K, 10K, or 30K MWCO).

The purified and concentrated DBCO-modified antibody and azide-modified siRNA were then mixed overnight at 4°C.

*The diagrams were created using Adobe Illustrator and PyMOL (Schrödinger)*.

---

**Figure S1.**
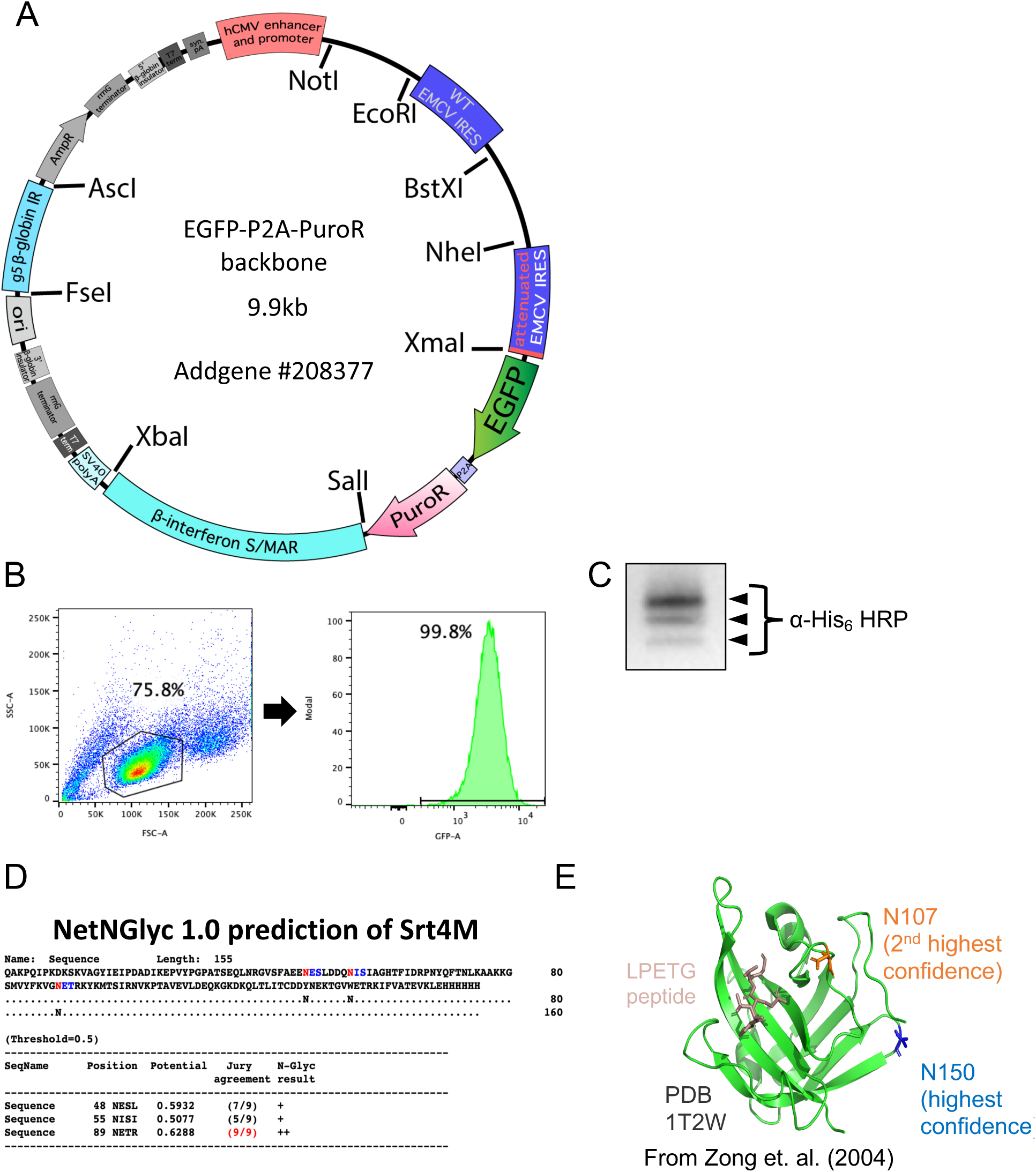
EGFP-P2A-PuroR vector backbone and characterization of Srt4M. **A.** EGFP-P2A-PuroR plasmid vector backbone with relevant unique restriction sites indicated. **B.** Gating strategy used to generate histograms of GFP expression intensity. **C.** Western blot staining for His_6_ tag from purified Srt4M using HRP anti-His tag antibody (BioLegend 652503) developed using ChemiDoc with chemiluminescent substrate (BioRad) **D.** Output from NetNGlyc - 1.0 prediction^53^ of N-glycosylation sites in Srt4M **E.** Sortase A crystal structure (PDB: 1T2W, Zong et al 2004^54^) with two highest confidence predicted glycosylation sites mapped in orange (N107) and blue (N150). The LPETG peptide is also shown in clam shell color. Image generated using PyMOL (Schrödinger).

**Figure S2.**
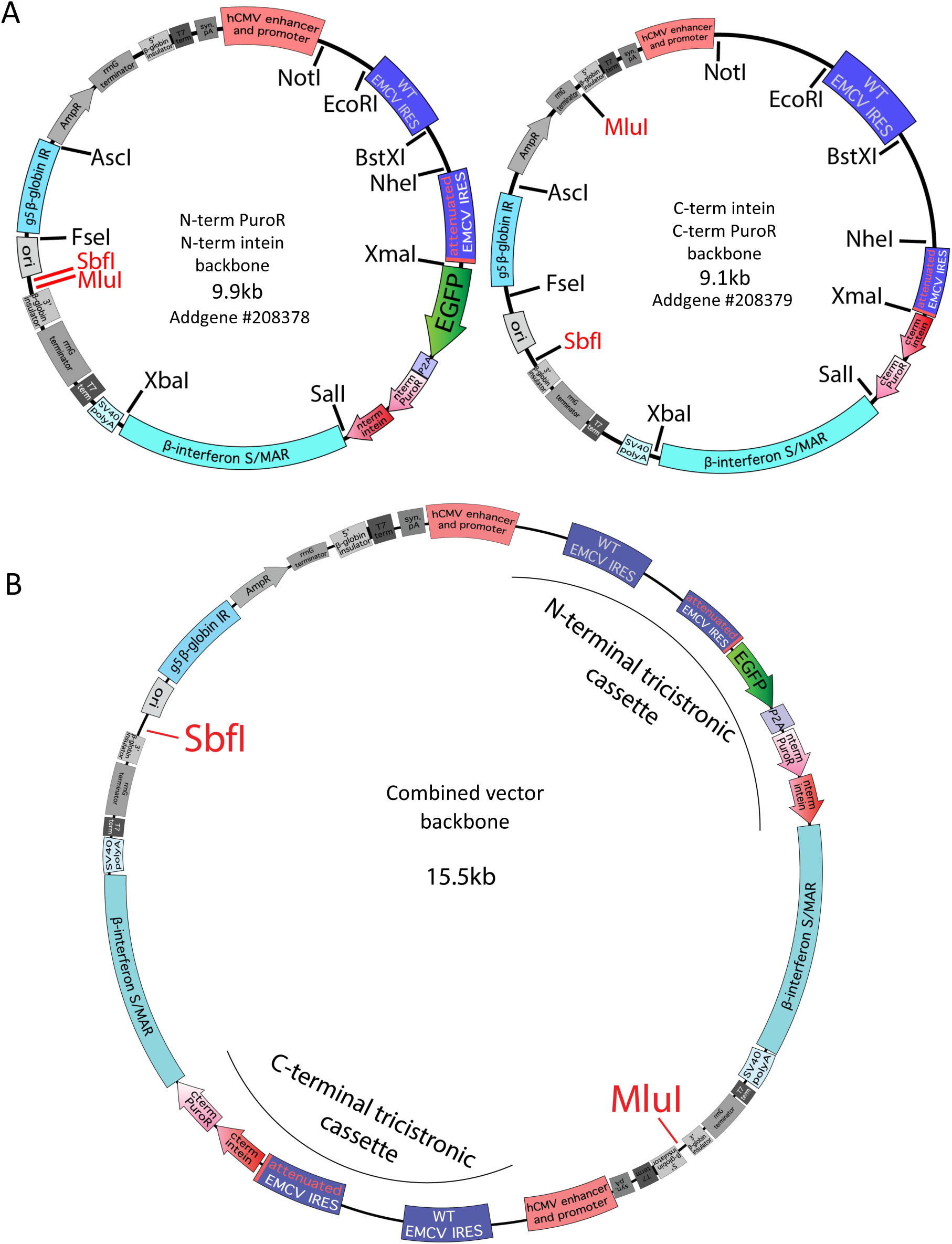
Split intein vector backbone maps. **A.** N-terminal PuroR N-terminal intein vector backbone with relevant unique restriction sites indicated (left). C-terminal intein C-terminal PuroR vector backbone with relevant unique restriction sites (right). SbfI and MluI sites are shown in red to highlight the different locations between the N-terminal and C-terminal plasmids whereas all other restriction enzyme sites are in identical locations in both plasmids. **B.** A combined vector containing both tricistronic cassettes linked to N- and C-terminal intein-puromycin cistrons.

**Table S1.**
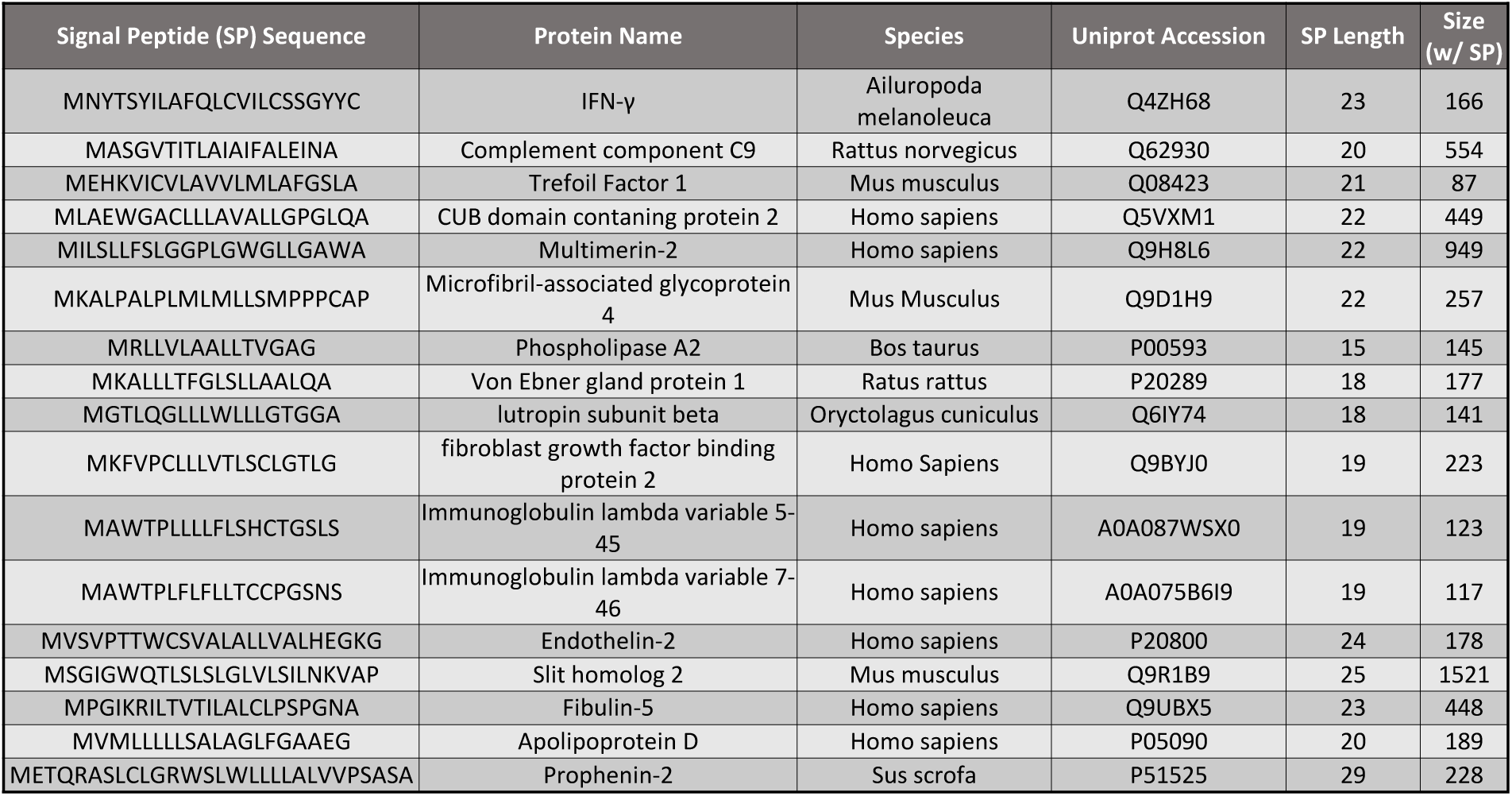
Signal peptide library of all peptides used that had predicted +1/+2 QA cleavage residues matching those of Srt4M Δ59.

**Table S2.**
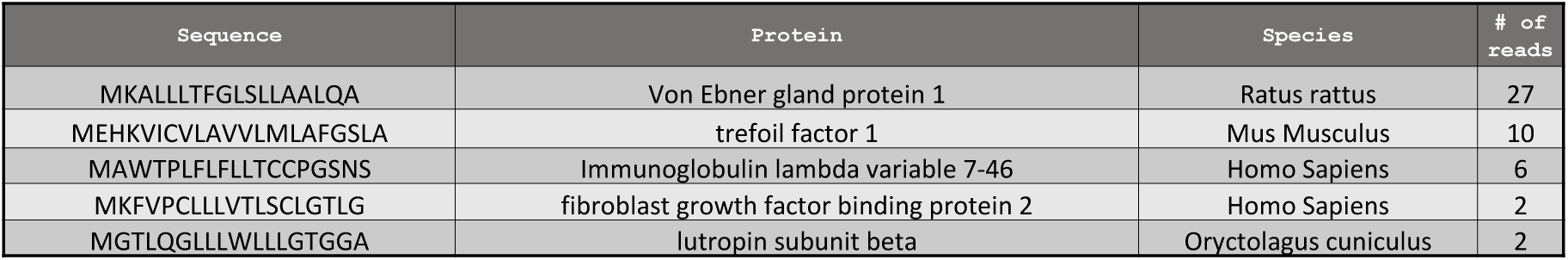
Signal peptides that were found after nanopore sequencing of PCR amplified DNA from cells selected with 20 ug/mL of puromycin for over 5 months.

